# A Master-Key DNA System Enabling Programmable Cross-Talks in Biomimetic Networks via An Artificial Chaperone

**DOI:** 10.64898/2026.06.16.732602

**Authors:** Wancheng Zhang, Minako Saito, Kenta Fujii, Naohiko Shimada, Atsushi Maruyama

## Abstract

Biological systems operate through complex molecular networks programmed by genetic information; however, constructing artificial systems with multilayered control remains a significant challenge. Here, we report a simple and integrated master-key system governed by lock DNA and master key DNA, reversibly switching diverse downstream processes ON/OFF and achieving dynamic cross-talks among distinct molecular components. The system utilizes cationic copolymer chaperones as control nodes, based on poly(L-lysine) or poly(allylamine) grafted with hydrophilic side chains, with a peptide nucleic acid (PNA) plug-in that grants sequence-specificity. We demonstrated two proof-of-concept systems: a nucleic acid-based catalytic network responsive to microRNA let-7b and a peptide-mediated transformation of lipid bilayers from two-dimensional sheets to three-dimensional vesicles. Both systems exhibited precise, modular, and programmable control with high robustness, mimicking the governing role of nucleic acids in biological systems. This strategy provides a versatile design framework for constructing biomimetic molecular networks and studying biological systems.

## Introduction

Biological systems operate through intricate molecular networks composed of interconnected biochemical reactions and regulatory loops. According to the central dogma, genetic information encoded in nucleic acid sequences serves as the master command of the production of functional molecules, thereby directing biological processes in a hierarchical and dynamic manner^1^. Through transcription and translation, nucleic acids and proteins are generated to perform particular functions, including enzymatic catalysis, energy production, membrane dynamics, and signaling transduction. These processes are further modulated by environmental factors such as pH ^2,3^, temperature^4^ and ionic conditions^5^, further synergistically enabling highly coordinated activities across cells, tissues and organs.

Importantly, certain biomolecules act as master regulators to control multiple downstream processes. For example, the master stem cell transcription factors such as Oct4, Sox2 and Nanog, are able to bind to specific DNA sequences and consequently regulating the expression of genes that control pluripotency, self-renewal and differentiation in stem cells ^6^. Such multilayered regulation, involving extensive and branched cross-talks among nucleic acids, proteins and lipids, weaves precise and highly efficient networks, systematically supporting the activities of living organisms.

Inspired by these natural systems, significant efforts have been devoted to constructing artificial systems capable of programmable control. Representative examples include CRISPR–Cas9 genome editing systems^7^, synthetic riboswitches^8,9^, and DNA-based signaling networks^10^, which enable sequence-specific regulation of molecular processes.

In addition, biomimetic systems integrating nucleic acid circuits with membrane components have been developed to achieve functions such as signal transduction, feedback regulation, and molecular computation^11,12^. These advances have found potential in fields of disease diagonsis^13^, medical therapies ^14^ and artificial cells^15,16^.

Despite these developments, constructing artificial systems that achieve multilayered regulation and dynamic cross-talk across different classes of biomolecules remains a major challenge. In particular, it is difficult to design a unified control system in which a single molecular input can orchestrate diverse downstream processes in a programmable and reversible manner. The complexity of integrating nucleic acids, polymers, peptides, and lipid assemblies into a coherent functional network has limited the development of such systems.

To address this challenge, we herein present a programable master key system that enables sequence-specific ON/OFF control of multiple molecular processes using a pair of DNA input. This system is based on a cationic copolymer chaperone, composed of poly-L-lysine (PLL) or poly(allylamine) (PAA) grafted with hydrophilic side chains of dextran (Dex) or polyethylene glycol (PEG), which enhances the folding and function of anionic biomolecules. To grant sequence-specific control, a peptide nucleic acid (PNA) module is covalently introduced as a plug-in, enabling hybridization-based regulation of the chaperone activity.

In this design, as illustrated in Figure 1, the copolymer acts as a functional “control node” that switches between active (ON) and inactive (OFF) states. Hybridization with a complementary lock DNA (L-DNA) suppresses the chaperone function, whereas subsequent strand displacement by a master key DNA (MK-DNA) restores activity and thus achieves high-performance, self-assembling nanodevices that are triggered by their own components (called private keys), ranging from DNA strand exchange reactions ^17,18^, DNAzymes ^19^, DNA circuits ^20,21^ to peptide-mediated membrane transformation^22,23^ according to our previous reports. This mechanism allows reversible and programmable control of downstream systems through DNA sequence inputs.

**Figure 1.**
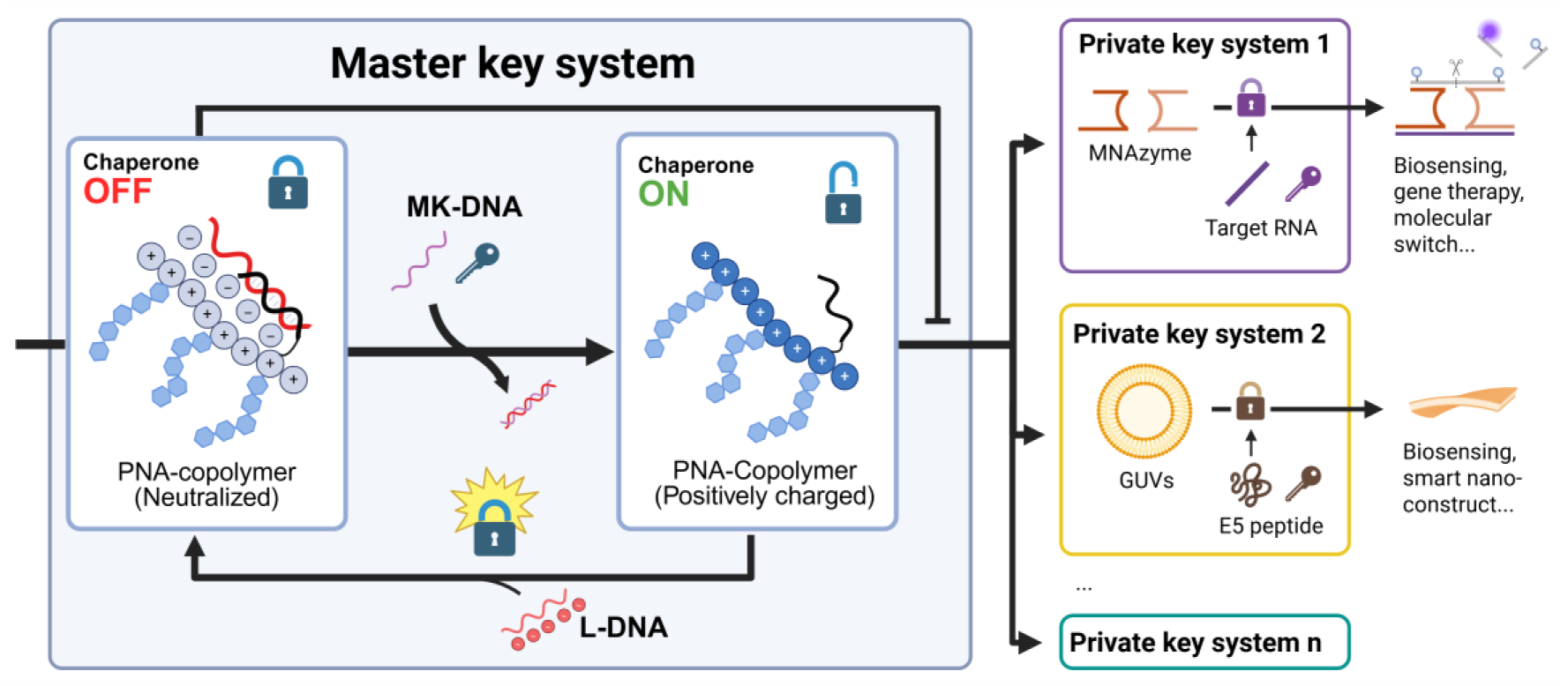
Scheme of the control chain of the master key-private key system through nucleic acid-responsive control of the chaperone function of the PNA-copolymer conjugate.

We demonstrate the versatility of this concept using two representative systems: (i) a nucleic acid-based catalytic network (MNAzyme) responsive to microRNA and (ii) a peptide-mediated transformation of lipid bilayers from two-dimensional nanosheets to three-dimensional vesicles. Through these systems, we establish a direct link between nucleic acid information and higher-order physicochemical transformations across multiple classes of biomolecules.

This master key architecture provides a generalizable framework for constructing integrated biomimetic networks with programmable cross-talks. By bridging nucleic acid signals with polymer-mediated control of peptide and membrane systems, this strategy provides a versatile building block for designing adaptive molecular systems in cells or artificial networks that mimic and study the processes in natural organisms, thereby expanding opportunities for applications in biosensing, smart therapeutics, and synthetic biology.

## Results and discussion

### 1. Master key system of PNA-copolymers controlling MNAzyme through DNA sequences

Figure 2a illustrates the operating principle of MNAzyme-based nanodevice targeting microRNA let-7b. Upon hybridization of the target let-7b with Partzyme A and Partzyme B, conformational rearrangement activates the catalytic core, leading to cleavage of the substrate strand ^24^. The RNA substrate is labeled with fluorophore-quencher pair on either side of the cleavage site, enabling fluorescence recovery upon cleavage and thus signaling the presence of the target sequence and reflecting the reaction rate.

**Figure 2.**
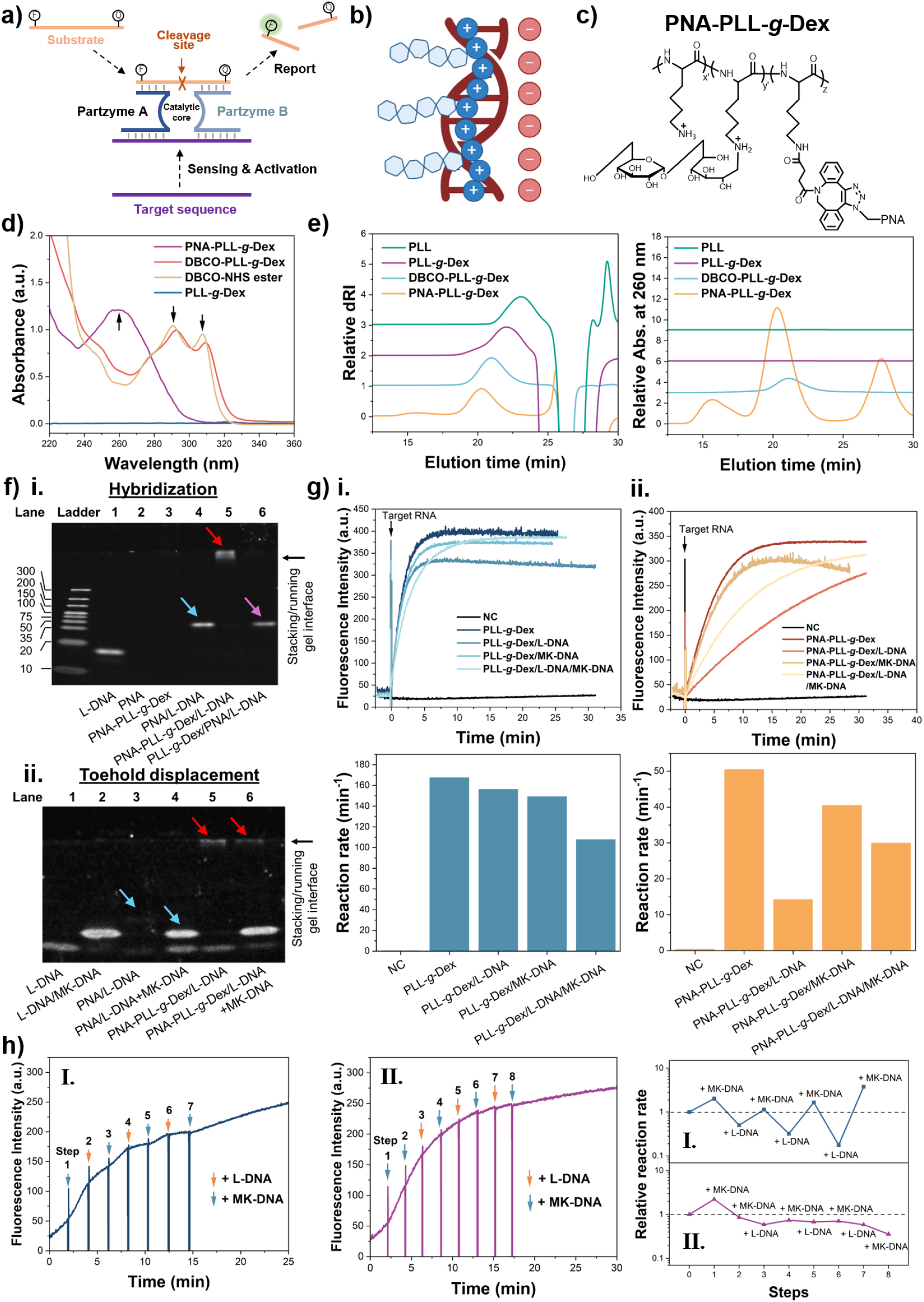
Construction of master key system for the control of MNAzyme and cross-talks among nucleic acid strands within. a) Scheme of principle of MNAzyme-based nanodevice. b) Scheme of binding between DNAs and cationic copolymers. c) Chemical structure of PNA-PLL*-g-*Dex. d) UV-Vis spectra of different copolymers and reagents during synthesis. e) GPC profiles of the copolymers by dRI (left) and UV detector (right), respectively. f) Performance of PNA-PLL*-g-*Dex in DNA strand exchange reactions, including i. hybridization and ii. toehold-mediated strand displacement, as evaluated by native PAGE. Red, blue and purple arrows point out groups with PNA-PLL*-g-*Dex, PNA without copolymers and copolymers and PNA not conjugated, respectively. g) Fluorescence change of MNAzyme system in presence of i. PNA-free copolymers and ii. PNA-PLL*-g-*Dex treated with different DNA strands in time-course (up) and corresponding reaction rates (down), respectively. h) Validations of reversibility and specificity of the master key system through multiple in-feed of MK-DNA and L-DNA by different orders (left and middle). Their relative reaction rates (right) were determined as the ratio of the reaction rate of each step to that of the previous step, with the relative reaction rates of the first step set to 1.

We previously reported that copolymer PLL-*graft*-Dex (PLL*-g-*Dex) reduces the electrostatic repulsion among nucleic acid strands upon complexation, thereby augmenting both hybridization efficiency and catalytic turnover (Figure 2b) ^25^. Consequently, the MNAzyme activity was significantly increased by up to three orders of magnitude compared to the unassisted system. However, this chaperone effect lacks sequence specificity using the prototype PLL*-g-*Dex, limiting its controllability.

To introduce sequence-programmable control, we designed a PNA-modified copolymer (PNA-PLL-*g*-Dex; Figure 2c), in which the PNA segment serves as a sequence-recognition module. The system was configured such that hybridization with lock DNA (L-DNA) suppresses the chaperone activity, whereas strand displacement by master key DNA (MK-DNA) restores it. In detail, the DNA sequence of microRNA miR-21, which has been wildly used as the marker of various diseases ^26^, was used as MK-DNA (22 nt), and its complementary sequences as L-DNA. The PNA was designed as the 13 nt sequence from 5’ terminus of miR-21.

Successful synthesis of PNA–PLL-g-Dex was achieved through strain-promoted azide–alkyne cycloaddition for its high orthogonality, high yield, limited side reactions, and simple operation^27–31^. PLL*-g-*Dex was first functionalized with dibenzocyclooctyne (DBCO) groups using DBCO-NHS ester (Scheme S1, Supporting Information). These intermediate products were named DBCO-PLL*-g-*Dex. The introduction of DBCO was verified by 1H-NMR, which showed broad aromatic proton signals of DBCO between 7.70-6.60 ppm, as compared to that of DBCO-NHS ester (δ (d6-DMSO) = 7.91-7.05) in Figure S1 (Supporting Information), which can be used to calculate the conjugation ratio of DBCO^32,33^. In UV-Vis spectra, the characteristic DBCO peaks also appeared at 293 and 310 nm (Figure 2d) ^34^. Upon the subsequent conjugation with azide-modified PNA, called azide-PNA (Table S1, Scheme S2, Supporting Information), gel permeation chromatography (GPC) profiles showed significantly elevated UV absorption at 260 nm at the copolymer fraction compared to DBCO-PLL*-g-*Dex under normalized dRI peak of the copolymers (Figure 2e), indicating presence of conjugated PNA. The forward shift of the copolymer peak further suggested an increase in the molecular weight. Consistently, UV-Vis spectra of the purified product showed PNA absorption at 260 nm together with reduced DBCO peaks (Figure 2d), supporting successful PNA incorporation. For the nomenclature, PLL*-g-*Dex, DBCO-PLL*-g-*Dex and PNA-PLL*-g-*Dex were named as 8kXD, 8kXDYDBCO and 8kXD-ZPNA, respectively, as 8k represents the average molecular weight of the PLL main chain of 8,000, X means the wt% of the dextran side chain, Y means the mol% of introduced DBCO moiety and Z means the mol% of conjugated PNA strands. The detailed compositions of the copolymers are listed in Table S2 (Supporting Information). The final product contained 89.7 wt% of dextran and 3.3 mol% of PNA strands and thus named as 8k90D-3.3PNA. This PNA content corresponds to theoretical neutralization of 72.6 % of cationic charges upon hybridization with L-DNA ideally, sufficient for effective copolymer shielding.

The performance of hybridization and toehold displacement of the PNA copolymers were evaluated by native polyacrylamide gel electrophoresis (Native PAGE). As shown in Figure 2f-I, for PLL-g-Dex mixed with free PNA (lane 6), a band corresponding to PNA/L-DNA hybrids (pink arrow) appeared, indicating dissociation of noncovalently associated hybrids from the copolymer. In contrast, the PNA-PLL-g-Dex lane (lane 5) produced a bright band (red arrow) near the stacking/running gel interface, attributed to PNA-copolymer/L-DNA complexes that could not migrate due to their large size, low charge density. Such band was not observed with MK-DNA (Figure S2, Supporting Information). This confirmed stable L-DNA retention by PNA-PLL-g-Dex even in the presence of excess competitive polyanion polyvinylsulfonic acid (PVS). Upon addition of MK-DNA (Figure 2f-ii), a bright band of L-DNA/MK-DNA duplex appeared in lane 6, demonstrating effective toehold displacement. These results demonstrate that DNA sequence inputs can dynamically regulate the state of the copolymer.

We first examined whether DNA could sequence-specifically regulate the chaperone activity of PNA-copolymers. As verified in Figure 2g-i, in the absence of target RNA, PLL-g-Dex mixed with MNAzyme components and substrate at N/P = 1 (the ratio between concentrations of positively charged amine on the copolymer and negatively charged phosphate on MNAzyme and substrate combined) produced negligible fluorescence, whereas addition of target RNA switched on MNAzyme cleavage, confirming its role as the private key. The prototype PLL-g-Dex significantly accelerated the initial reaction rate (167.5 min^-1^) by 420-fold compared with the no-copolymer control (NC, 0.4 min^-1^), demonstrating strong chaperone-mediated activation. Addition of L-DNA, MK-DNA, or their hybrids with the same strand concentration only partially reduced this effect with reaction rates of 156.1 min^-1^, 149.1 min^-1^ and 107.6 min^-1^ respectively, indicating that negatively charged polymers could attenuate the chaperone function in a very limited manner and lacked efficient sequence specificity.

PNA-PLL*-g-*Dex (Figure 2g-ii) also resulted in 126-fold increase of the initial reaction rate (50.5 min^-1^) than NC group, although less strongly than PLL*-g-*Dex possibly due to steric hindrance from PNA strands. Importantly, pre-annealing PNA-PLL-g-Dex with L-DNA reduced the reaction rate by 3.54-fold to 14.2 min^-1^, whereas MK-DNA or L-DNA/MK-DNA mixtures caused much weaker inhibition by 1.2 and 1.7-fold respectively, confirming L-DNA-specific switch-off behavior.

Furthermore, sequential addition of L-DNA and MK-DNA resulted in periodic modulation of reaction rates (Figure 2h-I) in a fast switch manner (Figure S3, Supporting Information), demonstrating dynamic and reversible control. With excess MK-DNA maintained the copolymer in an unlocked state and suppressed oscillation (Figure 2h-II). These results establish that the master key–private key architecture enables programmable regulation of nucleic acid catalysis with high specificity and quick responses.

### 2. DNAs mastering peptide-mediated lipid bilayer conversion through PNA-modified PAA copolymers

We further extend the master key system to peptide–lipid systems. Exampled by E5 peptide, the copolymer PAA-*graft*-Dex (PAA-*g*-Dex) chaperones its folding into α-helix and cause GUV membrane fracture. The complexes assemble at the edge of the bilayer fragments to stabilize the bilayers as 2D nanosheet (Figure S4a, Supporting Information)^23^. Addition of competing polyanions such as PVS or DNA disassembled E5/PAA-g-Dex complexes triggering destabilization of bilayers and autonomous conversion back to 3D vesicles under confocal laser scan microscopy (CLSM) in Figure S4b (Supporting Information). Without E5, no bilayer conversion could be realized, confirming its key role. In addition, we have reported that by hydrophobically anchoring both palmitated E5 (E5-pal) and the stearylated copolymer (PAA*-g-*Dex-SA, Scheme S3, Supporting Information), the converting nanosheets have high robustness against the electrostatic interference of free polyanions in the environment ^35^.

To allow sequence-specific control of lipid membrane topology, PNA-modified PAA-*g*-Dex (Figure 3a) and PAA-*g*-Dex-SA were synthesized through SPAAC similar as above, named respectively as PNA-PAA*-g-*Dex and PNA-PAA*-g-*Dex-SA (Scheme S4-S5, Supporting Information). The products showed 1H-NMR profiles (Figure S5, Supporting Information) similar to those of the PLL-based copolymers above. GPC analysis revealed nearly complete consumption of azide-PNA and increased UV absorption at 260 nm (Figure 3b), while UV–Vis spectra showed an increased A260/A310 ratio with higher PNA feed (Figure S6a, Supporting Information), confirming PNA conjugation. The PNA conjugation ratio was calculated from PNA consumption in GPC profiles as above. The PAA*-g-*Dex, PAA*-g-*Dex-SA, DBCO-PAA*-g-*Dex, DBCO-PAA*-g-*Dex-SA, PNA-PAA*-g-*Dex and PNA-PAA*-g-*Dex-SA were respectively coded as 5kXD, 5kXDYS, 5kXDZDBCO, 5kXDYSZDBCO, 5kXD-WPNA and 5kXDYS-WPNA, where 5k denotes the PAA backbone molecular weight, X the dextran wt%, Y the SA mol%, Z the DBCO mol%, and W the PNA mol% relative to monomer contents respectively. The compositions of the products are listed in Table S2 (Supporting Information). Native PAGE further confirmed that these PNA-copolymers retained hybridization and toehold displacement abilities (Figure S6b, Supporting Information), supporting DNA-mediated regulation of copolymer function.

**Figure 3.**
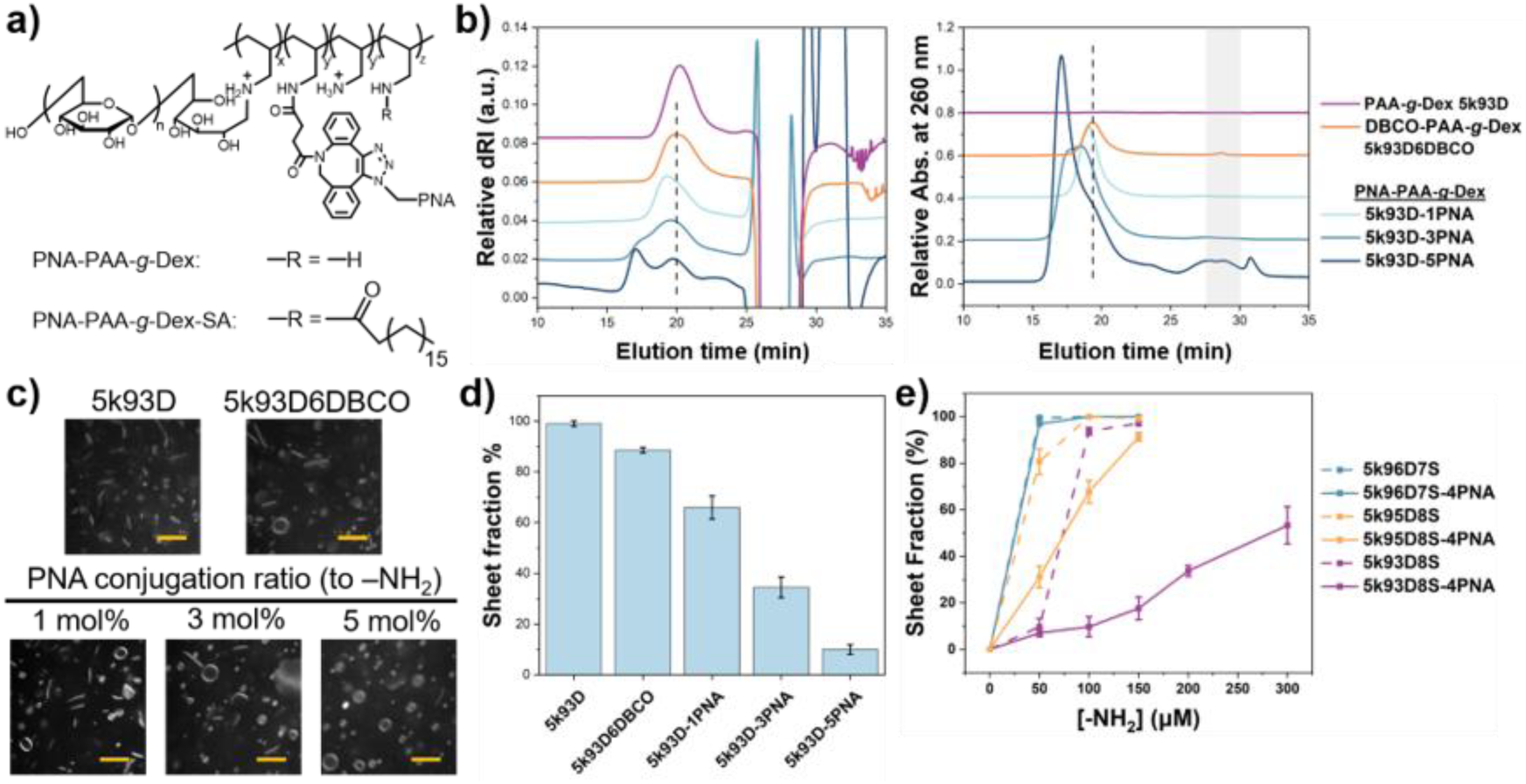
Synthesis and activity assessment of PNA-copolymer in membrane conversion. a) Chemical structures of PNA-PAA*-g-*Dex and PNA-PAA*-g-*Dex-SA. b) Comparison of GPC profiles of 5k93D, 5k93D6DBCO and the PNA-copolymer reacted from it after click chemistry with different PNA contents through dRI (left) and UV detectors (right) respectively. Grey zone represents where residual PNA eluted. c) Examine of performance of sheet formation before and after DBCO and PNA modification with the fixed concentration of 500 μM lipid, 10 μM E5 and 100 μM copolymers using CLSM, respectively, and d) their corresponding sheet fractions quantified. Scale bar: 5 μm. f) Sheet fractions yielded by different PAA*-g-*Dex-SA and their PNA conjugates, with fixed lipid concentration of 500 μM and E5-pal of 7.5 μM.

However, GPC profiles of PNA-copolymers showed that increasing PNA content resulted in the emergence of higher-molecular-weight fractions with higher UV absorption (Figure 3b), suggesting aggregation of the copolymers with more grafted PND strands. This behavior is attributed to the relatively hydrophobic backbone of PNA ^36^, which promotes intermolecular association. This aggregation hindered the copolymer function, presumably by disturbing complexation of the copolymers with E5 peptide and further organization at the bilayer edge to form stable nanosheets. At fixed 93 wt% dextran, increasing PNA content from 1 to 5 mol% reduced the sheet fraction from approximately 100% to approximately 10.0% with the same cation concentration of the copolymer, whereas PAA-g-Dex and DBCO-PAA-g-Dex retained good nanosheet-forming ability (Figure 3c and 3d). These results highlight a critical trade-off between sequence programmability and structural integrity.

To address this issue, the composition of the copolymer was optimized by increasing dextran content to enhance solubility and suppress aggregation (Figure S7, Supporting Information), noted by the decreased of early-eluted fraction, although excessive dextran may introduce steric hindrance ^37^. The sheet fraction increased from 7.0 ± 1.5% to ∼100% with the increase of dextran content from 93 to 96 wt% (Figure 3e and S8, Supporting Information). Balancing solubility, PNA content, and nanosheet formation, 5k96D8S-4PNA, which yielded nearly 100% sheet fraction, was selected for further evaluation.

Using the optimized PNA–PAA-g-Dex system, we investigated DNA-responsive control of membrane morphology. As schemed in Figure 4a, nanosheets formed by peptide–copolymer assembly undergo transformation into vesicles upon disruption of the chaperone function. Nanosheets formed with 5k96D8S-4PNA were treated with same amount of L-DNA and MK-DNA, respectively, at [A]/[C] = 0.5, where [A] is the concentration of the DNA anions and [C] is the concentration of the copolymer cations, respectively. Under nanomolar DNA conditions relevant to practical applications (147 nM), lipid membranes treated with non-complementary MK-DNA showed only limited conversion to vesicles, whereas L-DNA achieved significant sheet-to-vesicle transformation, yielding approximately 80% vesicles (Figure 4b, c). This strong sequence specificity was attributed to base-pairing interactions between PNA and L-DNA, which dominate over nonspecific electrostatic interactions.

**Figure 4.**
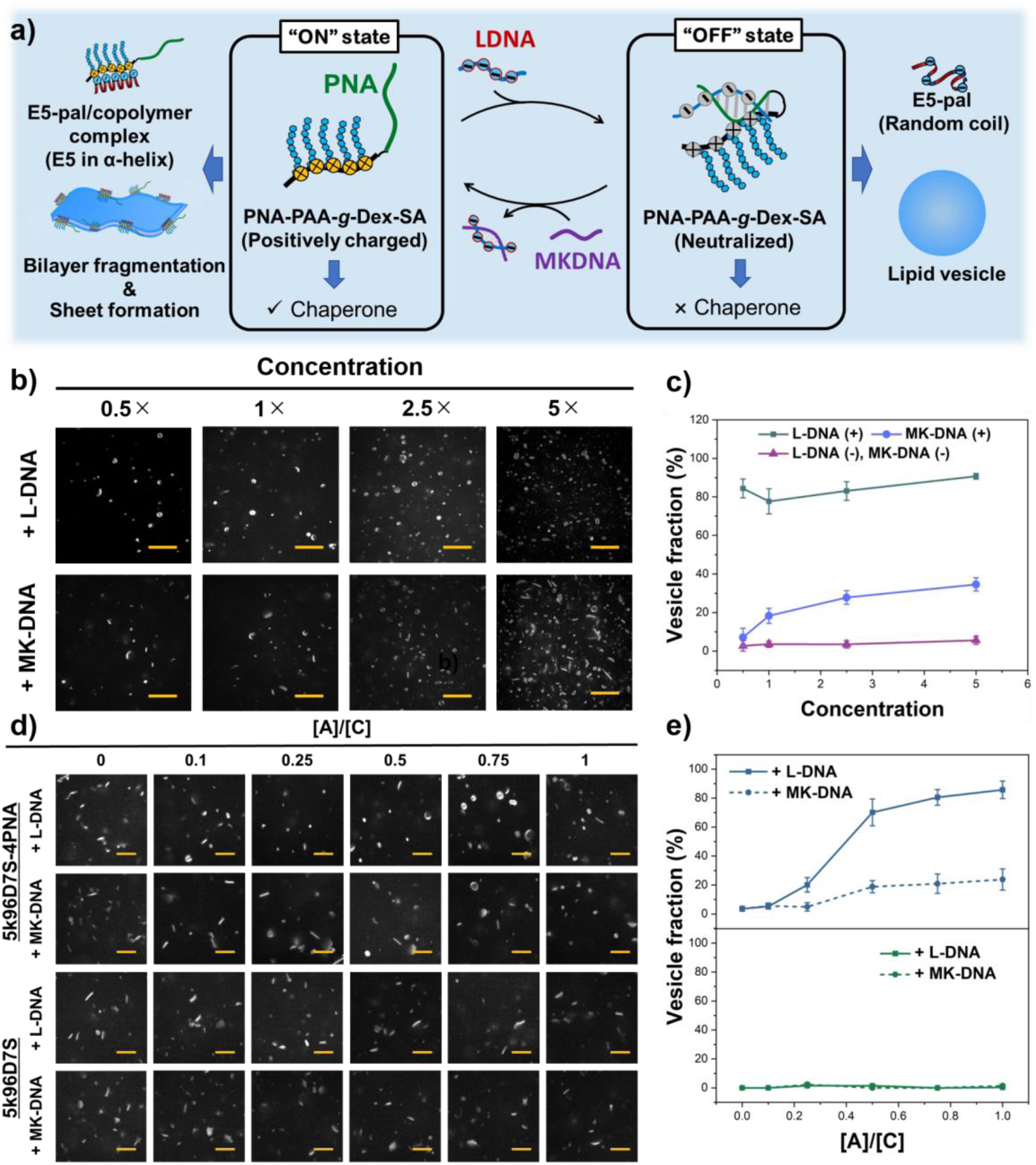
Membrane conversion achieved through cross-talks among DNA, peptide, lipid and PNA-copolymer chaperone. a) Scheme of master key system for the control of lipid membrane structures through specific DNA sequences. b) Influence of overall concentration on lipid membranes (60 μM) in presence of 0.9 μM E5-pal and 6 μM 5k96D7S-4PNA after feeding L-DNA or MK-DNA ([A]/[C] = 0.5) using CLSM respectively, and c) the corresponding vesicle fractions counted. Scale bar: 5 μm. d) Sheet-to-vesicle conversion of lipid membranes formed with 5k96D7S-4PNA and 5k96D7S by DNAs at different [A]/[C] ratio ([Lipid] = 60 μM, [copolymer] = 6 μM, [PNA] = 0.24 μM, [E5-pal] = 0.9 μM. Scale bar: 5 μm) and e) the quantification of vesicle fraction.

Such specificity persisted with varied overall concentration of the whole solution, although MK-DNA induced more nonspecific transformation at higher concentrations, likely due to enhanced nonspecific interactions (Figure 4b, c and S9, Supporting Information), while the nanosheets preserved when not treated with DNAs (Figure 4d, e). These results demonstrate that sequence-specific control is robust using the dual-anchoring system under complex conditions.

We next examined the influence of DNA length on system performance. Longer L-DNAs of 30 and 40 nt by adding adenines to the 5′ terminus and named 8A+L-DNA and 18A+L-DNA, respectively, were used to mimic longer control strands in practical applications. All L-DNAs followed sigmoidal dependence of sheet-to-vesicle conversion (Figure 5a–c), likely because of a combined effects of less unoccupied PNA, decreased cationic attraction and increased anionic repulsion (Figure 5d). Longer L-DNA strands induced more effective nanosheet-to-vesicle transformation at lower concentrations due to higher charge density per strand, as the vesicle fraction increased from 85.7 ± 6.0% for 22 nt L-DNA to 94.0 ± 2.5% for 40 nt L-DNA at [A]/[C] = 1. These findings indicate that the system can be tuned through sequence design, providing additional flexibility for programming molecular responses.

**Figure 5.**
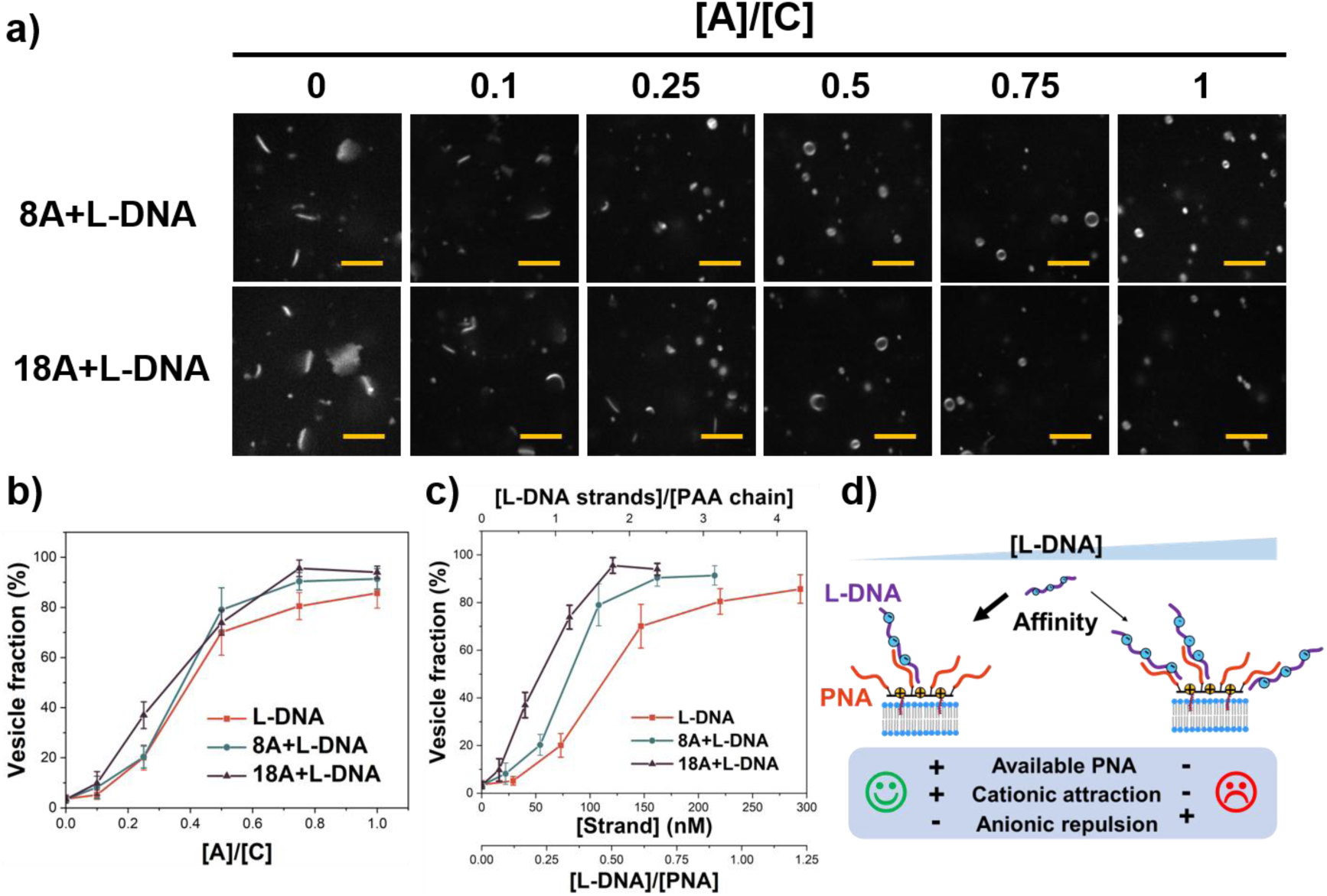
The same complementing motif overrides sequence context to dictate the controllability. a) Sheet-to**-**vesicle conversion of lipid membranes (60 μM) by L-DNAs with different lengths at various [A]/[C] ratio and the vesicle fractions were plotted in relation to: b) [A]/[C] ratio and c) [strand] (equivalent to [DNA strand] / [PAA chain] and [DNA]/[PNA]), respectively. [E5-pal] = 0.9 μM and [5k96D7S-4PNA] = 6 μM, incubation at 37 ℃. Scale bar: 5 µm. d) Sketch of proposed mechanism of S-like tendency of increase of vesicle fraction by L-DNAs.

Finally, we investigated the reversibility of the system by introducing MK-DNA to displace L-DNA. As shown in Figure 6a, addition of MK-DNA induced partial reversion of vesicles toward nanosheet-like structures. Increasing MK-DNA promoted partial re-transformation of vesicles into nanosheets, although many membrane structures became too small for reliable CLSM observation. Therefore, negative-stained transmission electron microscopy (nsTEM) was employed. Before and after the addition of MK-DNA, the membranes with regular contours, identified as vesicles, or those with irregular shapes both existed (Figure S10, Supporting Information) ^38,39^, and it revealed a reduction in membrane size following MK-DNA addition (Figure 6b), indicating reactivation of the copolymer–peptide system and further membrane fragmentation through MK-DNA-mediated displacement of L-DNA. These results demonstrate that membrane morphology can be reversibly regulated through DNA strand displacement. In other words, DNA information was transferred into lipid membrane topology through peptide and copolymer chaperones in a two-layered master key–private key system. This platform may support biosensing, controlled release, nanodevice assembly, and origin-of-life studies.

**Figure 6.**
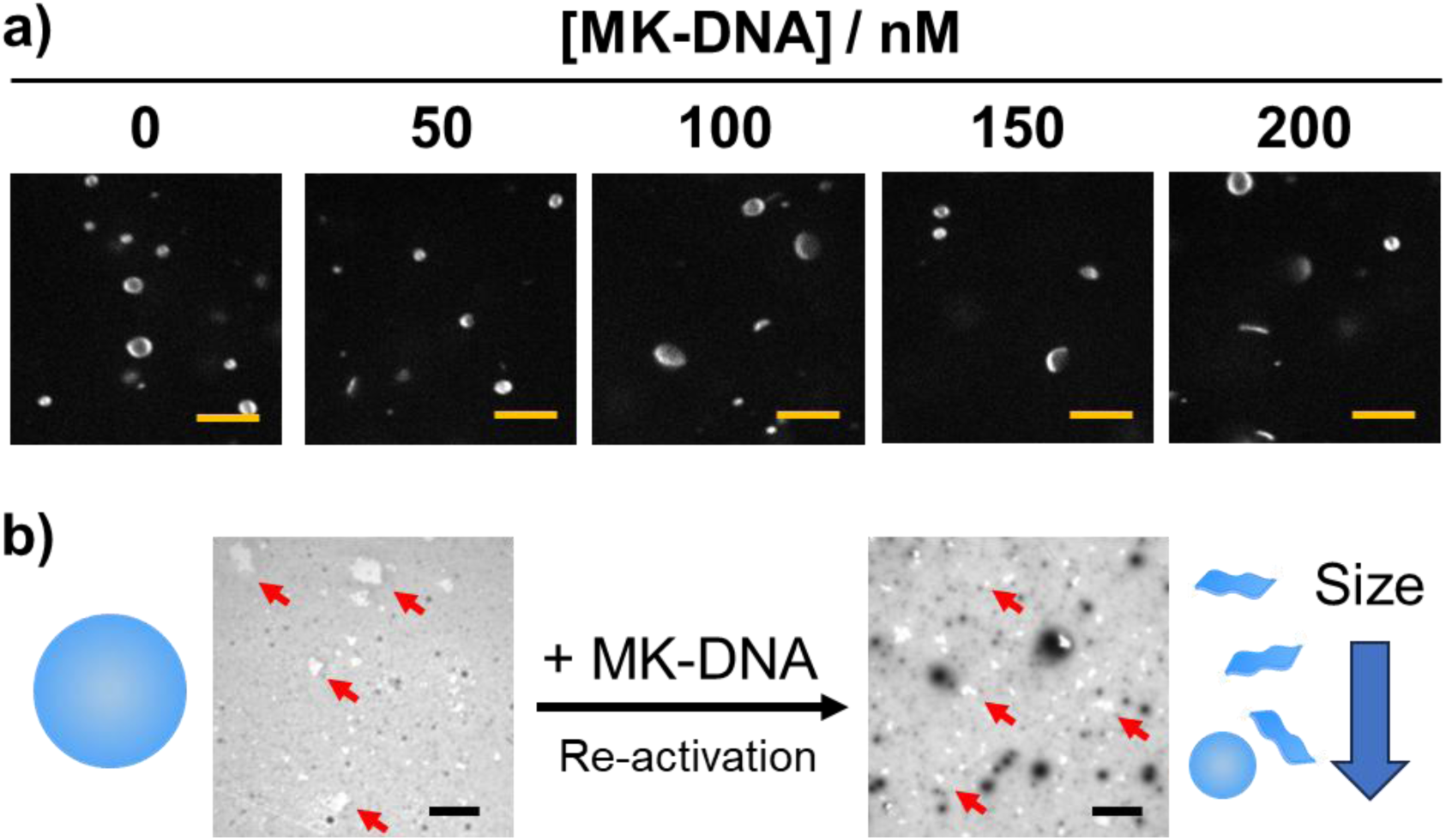
Reversed switch of the master0key system by MK-DNA. a) Re-conversion of vesicles to sheets by feeding different amount of MK-DNAs into the suspension with 56 μM lipids, 0.84 μM E5-pal and 5.6 μM 5k96D7S-4PNA, 137 nM of L-DNA ([Anion of L-DNA]/[Cation of copolymer] = 0.5), incubated at 37 ℃. Scale bar: 5 μm. b) Decrease of sizes of lipid membrane (red arrow pointed) after MK-DNA in-feed observed using TEM. Before TEM observation, 56 μM lipids were treated with 0.84 μM E5-pal and 5.6 μM 5k96D7S-4PNA, 137 nM of L-DNA ([Anion of L-DNA] / [Cation of copolymer] = 0.5) subsequentially. Scale bar: 10 µm.

## Conclusion

In this study, we have established a programmable master key–private key system that allows sequence-specific ON/OFF control of diverse molecular processes through a pair of DNA inputs. By integrating a cationic copolymer chaperone with a peptide nucleic acid (PNA)–based sequence-recognition plug-in, we created a molecular control node capable of dynamically regulating downstream activities in a reversible and highly specific manner. The generality of this concept was demonstrated using two distinct systems: a nucleic acid-based catalytic network (MNAzyme) and a peptide-mediated lipid membrane transformation system. In both cases, the chaperone function of the copolymer was precisely modulated through DNA hybridization and strand displacement, enabling programmable switching of enzymatic activity and membrane topology.

Importantly, this work establishes a direct link between nucleic acid information and higher-order physicochemical transformations across distinct classes of biomolecules. The ability to propagate sequence-defined signals through polymers to control peptides and lipid assemblies highlights a new design principle for integrating molecular functions into hierarchical systems. Given its modularity and versatility, this master key architecture provides a general framework for constructing biomimetic networks with programmable cross-talks. This strategy may boost the development of advanced molecular systems for applications in biosensing, smart therapeutics, and adaptive nanomaterials, while also offering a simplified platform for studying information flow and functional integration in modern and prebiotic biological systems.

## Acknowledgement

This work was financially supported by JSPS KAKENHI (4H00791 and 24K21246), a Grant-in-Aid by the Cooperative Research Program of “Network Joint Research Center for Materials and Devices”, the Promotion of Distinctive Joint Research Center Program (JPMXP 0621467946) and JST, the establishment of university fellowships towards the creation of science technology innovation (JPMJFS2112).

## Experimental section

### Materials

Poly-L-lysine hydrobromide (PLL·HBr) with average Mw of 8,000, sodium cyanoborohydride (NaBH_3_CN), 4-(2-hydroxyethyl)-1-piperazineethanesulfonic acid (HEPES), poly(vinylsulfonic acid, sodium salt) and dibenzocyclooctyne-N-hydroxysuccinimidyl ester (DBCO-NHS ester) were purchased from Sigma-Aldrich. Poly(allylamine) hydrochloride (PAA·HCl) aqueous solution with average Mw of 5,000 was provided by Nitto Boseki Co. It was precipitated in methanol at 4 ℃ and vacuum-dried overnight to obtain a white powder. Dextran (average Mw: 10,000) was purchased from Funakoshi Co. Ltd. The 1H-Benzotriazol-1-yloxy-tri(pyrrolidino)phosphonium hexafluorophosphate (PyBOP) was obtained from Watanabe, and N,N-diisopropylethylamine (DIPEA) was purchased from Nacalai Tesque, Inc. Stearic acid (SA) was acquired from TCI Co., Ltd. The E5 peptide, the sequence of which is H-GLFEAIAEFIEGGWEGLIEG-OH, and E5-pal, where E5 was conjugated with palmitic acid on the C terminus, were purchased from Genenet. The 1,2-dioleoyl-sn-glycero-3-phosphocholine (DOPC) and 1,2-dioleoyl-sn-glycero-3-phosphoethanolamine-N-(lissamine rhodamine B sulfonyl) (ammonium salt) (LR-DOPE) were acquired from Avanti Polar Lipids. N-(carbonyl-methoxypolyethyleneglycol)-1,2-distearoyl-sn-glycero-3-phosphoethanolamine (DSPE-PEG2k) was purchased from Yuka Sangyo Co. Azide activated PNA (azide-PNA) was provided by Biologica Co. The DNAs and RNAs were synthesized by Fasmac. The sequences of all involved nucleic acids are listed in Table S1. Other reagents used were purchased from FUJIFILM Wako. All reagents were used without further purification unless stated otherwise.

### Synthesis of PLL*-g-*Dex, PAA*-g-*Dex, PAA*-g-*Dex-SA, DBCO-PLL*-g-*Dex, DBCO-PAA*-g-*Dex and DBCO-PAA*-g-*Dex-SA

#### 1) PLL*-g-*Dex and PAA*-g-*Dex

PLL*-g-*Dex and PAA*-g-*Dex were synthesized by aldehyde-amine reaction as previously reported.^40^ Firstly, PLL·HBr or PAA·HCl was dissolved in 0.1 M sodium borate buffer (pH 8.5). Dextran and NaBH_3_CN were added in 3 aliquots over 3 d while stirred at 40 ℃. Unreacted dextran was removed using a cation exchange column (Toyopearl CM-650C), followed by eluting with a solution containing 1.0 M acetic acid and 0.5 M NaNO_3_ to collect the product. The copolymer was further dialyzed in MilliQ for 3 d then freeze-dried for another 3 d to obtain white powder.

#### 2) PAA*-g-*Dex-SA

PAA*-g-*Dex-SA was prepared by conjugating SA to the primary amines of PAA*-g-*Dex through PyBOP coupling (Scheme S2). In brief, PAA*-g-*Dex, PyBOP and SA stocks were dissolved in DMSO. To suppress side coupling reaction of SA with hydroxyl groups of dextran, ethanol was firstly added to PAA*-g-*Dex (final DMSO:Ethanol = 9:1). Then, PyBOP (0.1 eq.), SA (0.1 eq.) and DIPEA (0.3 eq.) were added to PAA*-g-*Dex (1 eq.) solution in order, and the mixture was incubated for 24 h at 40 ℃. The copolymer was purified by dialysis against DMSO then MilliQ using a regenerated cellulose membrane (MWCO: 3,500), followed by freeze-drying for 3 d to obtain a white powder. The SA grafting ratio was determined by analyzing PAA*-g-*Dex in d6-DMSO/D_2_O using ^1^H-NMR (400 MHz, Bruke) at 60 ℃.

#### 3) DBCO-PLL*-g-*Dex, DBCO-PAA*-g-*Dex and DBCO-PAA*-g-*Dex-SA

PLL*-g-*Dex, PAA*-g-*Dex or PAA*-g-*Dex-SA (0.01 mmol in-NH_2_, 1 eq.) was dissolved in HEPES buffer containing 10 mM HEPES and 140 mM NaCl (pH 7.4). DBCO-NHS ester (0.001 mmol, 0.1 eq.) dissolved in DMSO was added to the copolymer solution, followed by waterbed shaking for 24 h at room temperature. The final ratio of HEPES buffer and DMSO was 9:1. The unreacted DBCO was removed through washing by dichloromethane at 4 ℃ and dialyzed against MilliQ for 1 d at room temperature, and the mixture was freeze-dried for 3 d to obtain a white powder.

### Characterizations of the copolymers

The compositions of the PLL*-g-*Dex, PAA*-g-*Dex, PAA*-g-*Dex-SA, DBCO-PLL*-g-*Dex, DBCO-PAA*-g-*Dex and DBCO-PAA*-g-*Dex-SA were determined by 1H-NMR spectrometer (400 MHz, AVANCE II 400C, Bruker) at 60 ℃. PLL*-g-*Dex, PAA*-g-*Dex and DBCO-PLL*-g-*Dex were dissolved in D_2_O for the measurement, DBCO-PAA*-g-*Dex was dissolved in a mixture of d6-DMSO/D_2_O (v/v = 1:9), while PAA*-g-*Dex-SA and DBCO-PAA*-g-*Dex-SA were dissolved in d6-DMSO/D_2_O (v/v = 9:1). The concentration of all samples was fixed to 10 mg/mL.

The copolymers were analyzed using gel permeation chromatography (GPC, Jasco). The copolymers were dissolved respectively in MilliQ to obtain a solution of 10 mg/mL concentration and filtered through 0.45 μm membrane. One hundred microliter solution of each sample was injected into GPC instrument installed with Shodex SB804 and SB806M column, using a solution containing 0.5 M acetic acid and 0.5 M TEA (pH 7) as mobile phase. The elute was traced by RI (RI-4030, Jasco) and UV (UV-2070 Plus, Jasco) detectors at 260 nm.

UV-Vis absorption spectra of the PLL copolymer series dissolved in MilliQ were recorded using Nanodrop 1000 UV/Vis spectrophotometer (Thermofisher). Those of the PAA copolymer series dissolved in MilliQ were recorded with a UV-Vis spectrophotometer (UV-1650, Shimadzu).

### Synthesis of PNA-PLL*-g-*Dex, PNA-PAA*-g-*Dex and PNA-PAA*-g-*Dex-SA

PNA-PLL*-g-*Dex, PNA-PAA*-g-*Dex or PNA-PAA*-g-*Dex-SA was synthesized by mixing azide-PNA with DBCO-PLL*-g-*Dex, DBCO-PAA*-g-*Dex or DBCO-PAA*-g-*Dex-SA, respectively, in HEPES buffer (10 mM HEPES and 140 mM NaCl, pH 7.4), followed by thermal shaking for 24 h. Heating at 50 ℃ was employed to facilitate dissolution of azide-PNA under high-salt condition during reaction (Scheme S3).

The reaction mixture was purified and analyzed by GPC. Briefly, the reaction mixture was filter through 0.45 μm membrane and injected into GPC instrument. The mobile phase contained 0.5 M acetic acid and 0.5 M TEA (pH 7). The elute was analyzed by the detectors, and only the copolymer fraction was collected. The introduction ratio of PNA can be determined by subtracting the amount of unreacted azide-PNA from the total in-fed PNA, as no significant loss of PNA was detected during GPC measurement when comparing the quantity of in-fed PNA with that calculated from the peak area of PNA fraction in the UV profile using a extinction coefficient of 127,500 M^-1^ cm^-1^, as presented in Figure S11 and Table S3 (Supporting Information).

To obtain the powder of PNA-modified copolymer, ultracentrifuge filter unit (MWCO: 10k, Ultra-15, Amicon) was used to exchange the solvent of the collected copolymer to MilliQ. The product was freeze-dried for 1 d to yield a white powder then re-dissolved into HEPES buffer (10 mM HEPES and 140 mM NaCl, pH 7.4). The concentration of the solution of PNA-modified copolymer was determined through UV absorbance of PNA at 260 nm using Nanodrop or UV-Vis spectrophotometer.

### Assessment of hybridization and toehold displacement performance of PNA-modified copolymers by native-PAGE

For hybridization and specificity performances, PNA-PLL*-g-*Dex (or PNA-PAA*-g-*Dex), PNA along with PLL*-g-*Dex (or PAA*-g-*Dex), PNA alone and MK-DNA alone were respectively mixed with L-DNA in HEPES buffer (10 mM HEPES, 140 mM NaCl, pH 7.4) and heated at 90 ℃ for 5 min then cool to 25 ℃ in 2 h for annealing. The final mixture contained 2 μM L-DNA. L-DNA labeled with FITC were used during electrophoresis unless stated otherwise. For toehold displacement study, after annealing with unlabeled L-DNA, 2 μM MK-DNA (not labeled) was added to each mixture and incubated at room temperature for 30 min.

Before electrophoresis, the mixtures were treated with 10× excessive PVS (amount of monomers relative to that of the copolymers) to allow electrostatically bonded DNA to dissociate from the copolymers and migrate freely during electrophoresis.

Native PAGE gel composed of 20 % separation gel and 4.5 % stacking gel was prepared. The samples were mixed with 2× TrackIt Cyan/Yellow Loading Buffer (ThermoFisher) by half, and 8 μL of the mixtures was loaded into each well. The electrophoresis was run at 5 ℃ for 90 min at 100 V. The gels with FITC-L-DNA were imaged with a gel imager (LAS-3000, Fujifilm), while those with unlabeled L-DNA were imaged after staining by SYBR Gold for 30 min at RT.

### Fluorescence analysis of DNAzyme activity

The DNAzyme components, Mz PartA and Mz PartB was mixed with FQ substrates in HEPES buffer containing 10 mM HEPES, 150 mM NaCl, 0.1% Tween-20, pH7.3. The DNAs and copolymers were annealed and then added to yield a mixture with [Mz PartA] = [Mz PartB] = 2 nM, [FQ substrate] = 50 nM, [PNA]/[DNA] = 1 and N/P = 1. The fluorescence was recorded by fluorospectroscopy (JASCO FP-6500) at 37 °C under continuous stirring, to ensure the solution mixed well with subsequently added materials. To trigger the activation of DNAzyme, 5 mM of MnCl2 solution was injected into the mixture through the injection hole, followed by the in-feed of 1 nM target RNA. Reaction rates were derived from the slope at the very beginning of the fluorescence-time plot.

### Preparation of giant unilamellar vesicles (GUVs)

GUVs were prepared using a modified reverse phase method as previously reported ^41^. In brief, DOPC in diethyl ether, DSPE-PEG2k in chloroform and LR-DOPE in chloroform were mixed to yield a 500 μL solution with total phospholipids of 1.2 mM composed of DOPC: DSPE-PEG2k: LR-DOPE = 98.4:1.4:0.2 in mole. Another 500 μL of aqueous inner buffer (pH 7.4) containing 10 mM HEPES, 90 mM NaCl and 100 mM sucrose was added. The cocktail was vortex for 1 min, then sonicated for 4 min and finally vortex for another 1 min, followed by centrifugation at 20,000 × g for 2 min at 20 ℃. After removing the upper organic phase, the GUV suspension was washed with 500 μL outer buffer (pH 7.4) containing 10 mM HEPES and 140 mM NaCl by centrifugating at 5,000 × g for 10 min at 20 ℃ and discarding the supernatant for 3 repeats. The GUVs were finally suspended in ∼ 50 μL outer buffer. The lipid concentration of which was determined using LabAssay ™ Phospholipid Kit following the manufacturer’s instruction. In brief, 10 μL of GUV suspension was pipetted to an Eppendorf tube of 2 mL and sonicated for 10 min. A 1.5 mL of LabAssay test solution was subsequently mixed with the suspension and incubated at 37 ℃ for 10 min. The UV-Vis absorbance of the reaction mixture was then measured by UV-Vis spectrometer, and the concentration of lipids was determined by refereeing to the calibration curve.

### Liposome conversion driven by nucleic acids

The conversion of lipid membranes in responsive to specific DNA sequences were investigated in a cascade of stage A, B and C as shown in Scheme S6. In stage A, the capability of sheet formation with E5-pal chaperoned by PNA-PAA*-g-*Dex-SA was tested. The suspension of GUVs was incubated with E5-pal and subsequently PNA-PAA*-g-*Dex-SA at 37 ℃ for 30 min each. In stage B, the mixture in stage A was diluted by various magnitudes, and different DNA sequences were added to investigate the shielding abilities of DNA and system specificity. Upon addition, the mixtures were incubated at 37 ℃ for 30 min. In stage C, MK-DNA was added to the mixture in stage B aiming at triggering toehold displacement, subsequent re-activation of the copolymer and resulting vesicle conversion. After adding MK-DNA, the mixture was incubated at 37 ℃ for 30 min.

### Observation of morphologies of lipid membranes

Morphological observation of lipid membranes was conducted using CLSM. Ten microliter suspension of lipid membranes was dropped on a cover glass set on the microscope (IX71, Olympus) installed with spinning-disk confocal scanning unit (CSU-X1, Yokogawa), an EMCCD camera (iXon+ 893, Andor) and a 100× objective lens (UPlanSApo NA 1.4, Olympus). Liposomes labeled with LR-DOPE were excited with the laser sources of 561 nm for observation.

The sheet or vesicle fractions were determined by counting their numbers in the confocal microscopic images (n = 5) and expressed as mean ± STD.

Meanwhile, negatively stained transmission electron microscopy (nsTEM) was employed to observe lipid membranes prepared in stage B and C. Five microliter of samples were dopped on the copper grid and settled for 1 min at room temperature for attachment. The excess of solution was absorbed by filter paper. The phosphotungstic acid solution, the pH of which was adjusted to 6 by 1 M KOH, was dropped on the grid and stained for 1 min. The excessive stain was removed by filter paper again, and the grid was air dried. Then the samples were imaged using a JEOL JEM-1400Plus microscope at a voltage of 100 kV.

