## Supporting Information for "A Master-Key DNA System Enabling Programmable Cross-Talks in Biomimetic Networks via An Artificial Chaperone"

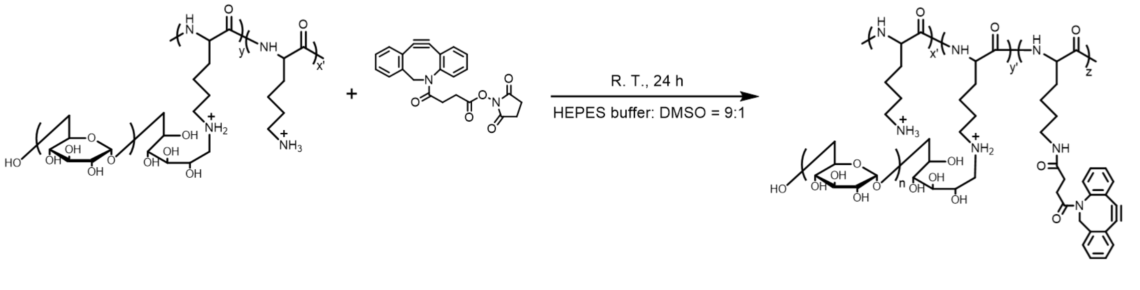


**Scheme S1.** Synthesis of DBCO-PLL-g-Dex.

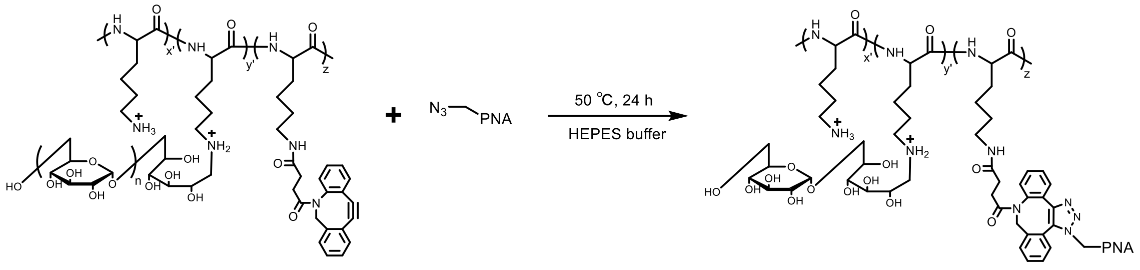


**Scheme S2.** Synthesis of PNA-PLL-g-Dex.

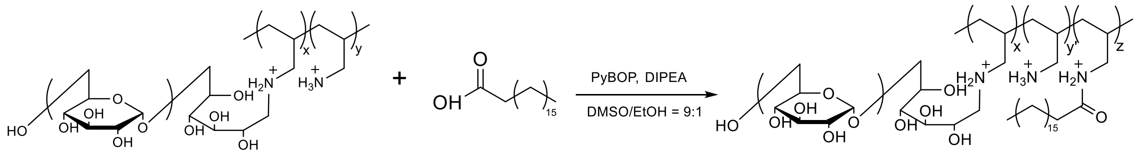


**Scheme S3.** Synthesis of PAA-g-Dex-SA.

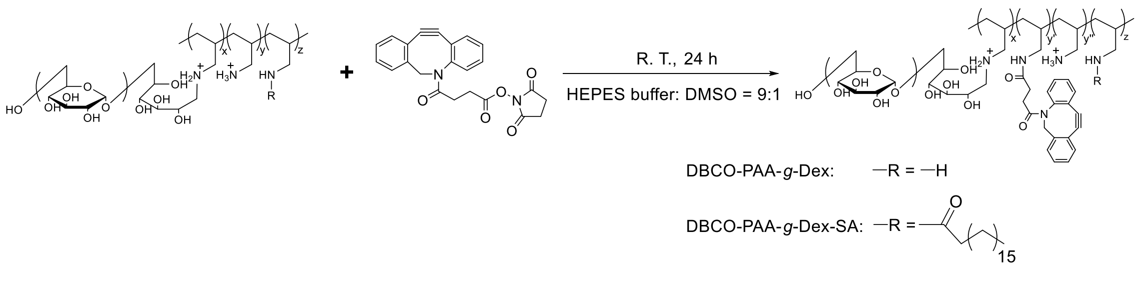


**Scheme S4.** Synthesis of DBCO-PAA-g-Dex or DBCO-PAA-g-Dex-SA.

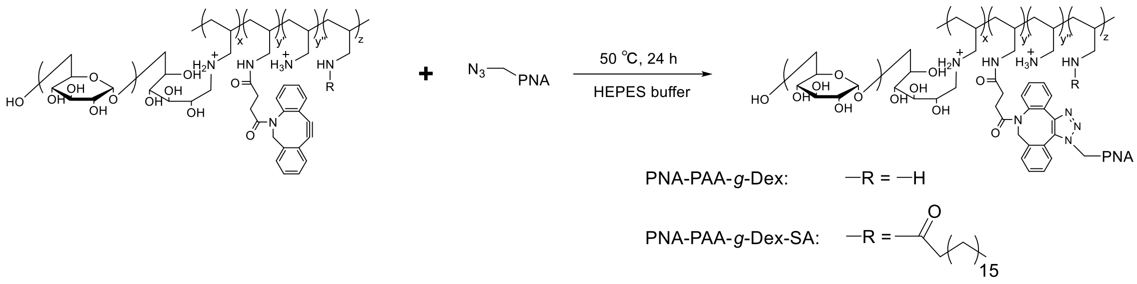


**Scheme S5.** Synthesis of PNA-PAA-g-Dex or PNA-PAA-g-Dex-SA.

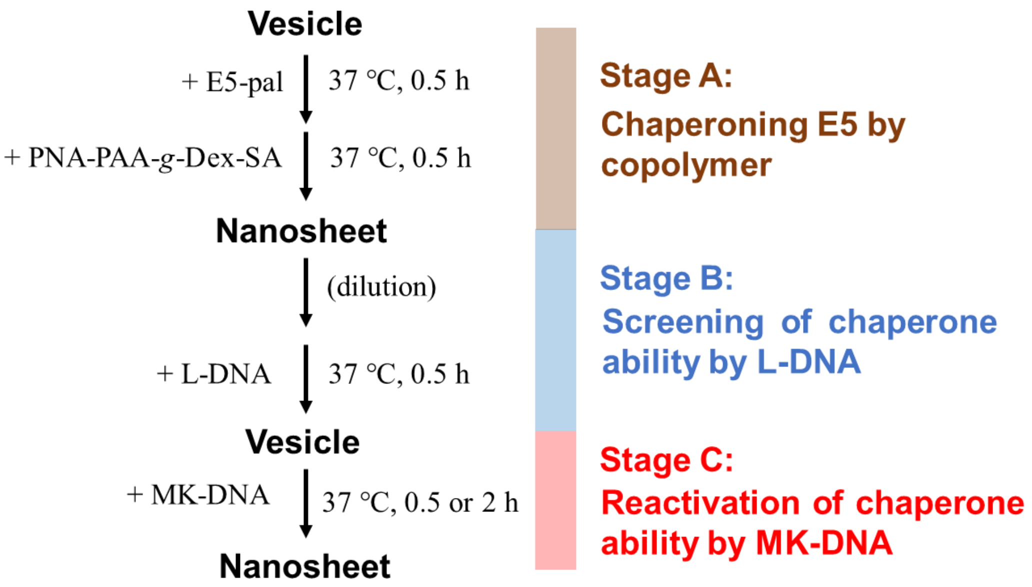


**Scheme S6.** Scheme of reversible conversion of lipid membranes controlled by DNA sequences.

**Table S1.** List of involved nucleic acid sequences.

| Code | Sequence | Length / nt |
|---|---|---|
| Mz PartA | 5’-CCAGGGAGGCTAGCTCTACTACCTCA-3’ | 26 |
| Mz PartB | 5’-AACCACACAACACAACGAGAGGAAAC-3’ | 26 |
| FQ substrate | 5’-GTTTCCTCguCCCTGG-3’ | 16 |
| Target RNA | 5’-UGAGGUAGUAGGUUGUGUGGUU-3’ | 22 |
| L-DNA | 5’-TCAACATCAGTCTGATAAGCTA-3’ | 22 |
| 8A+L-DNA | 5’-AAAAAAAATCAACATCAGTCTGATAAG CTA-3’ | 30 |
| 18A+L-DNA | 5’-AAAAAAAAAAAAAAAAAATCAACATC AGTCTGATAAGCTA-3’ | 40 |
| MK-DNA | 5‘-TAGCTTATCAGACTGATGTTGA-3’ | 22 |
| miR21-azide-PNA | *N*-Azide-TAGCTTATCAGAC-*C* | 13 |

**Table S2.** Chemical compositions of DBCO-PLL-g-Dex, DBCO-PAA-g-Dex and DBCO-PAA-g-Dex-SA.

| Copolymer type | Code | Dextran wt% | Dextran mol% | SA mol% | DBCO mol% | DBCO conjugation yield % |
|---|---|---|---|---|---|---|
| PLL-series | 8k90D7DBCO | 89.7 | 12.7 | - | 7.4 | 74.2 |
| PAA-series | 5k93D6DBCO | 93.2 | 7.8 | - | 6.4 | 63.9 |
| | 5k95D6DBCO | 94.7 | 10.2 | - | 6.5 | 64.7 |
| | 5k96D8DBCO | 96.1 | 14.0 | - | 8.1 | 81.0 |
| | 5k93D8S5DBCO | 93.2 | 7.8 | 7.7 | 4.7 | 46.6 |
| | 5k95D8S8DBCO | 94.7 | 10.2 | 7.5 | 8.4 | 84.2 |
| | 5k96D7S8DBCO | 96.1 | 14.0 | 7.0 | 8.5 | 84.7 |

**Table S3.** Comparison of amount calculated from PNA fraction in UV profiles of GPC and the amount of PNA in-feed.

| PNA calculated / nmol | PNA in-feed / nmol | Calculated / In-feed |
|---|---|---|
| 3.64 | 3.76 | 1.03 |
| 2.50 | 2.66 | 1.06 |
| 1.80 | 1.91 | 1.06 |

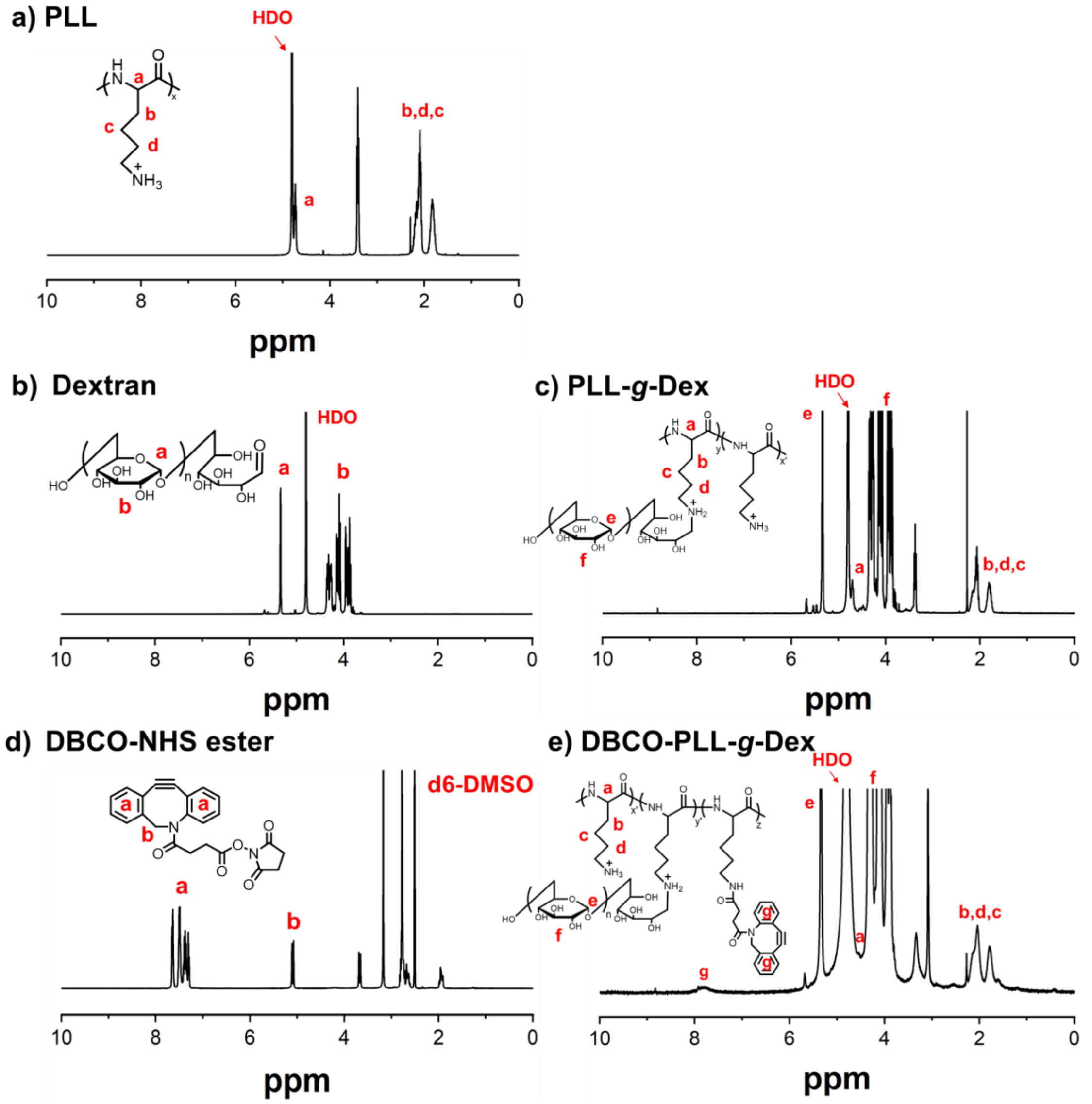


**Figure S1.** $^{1}$H-NMR spectra (400 MHz) of a) PLL, b) Dextran, c) PLL-g-Dex, d) DBCO-NHS ester and e) DBCO-PLL-g-Dex.

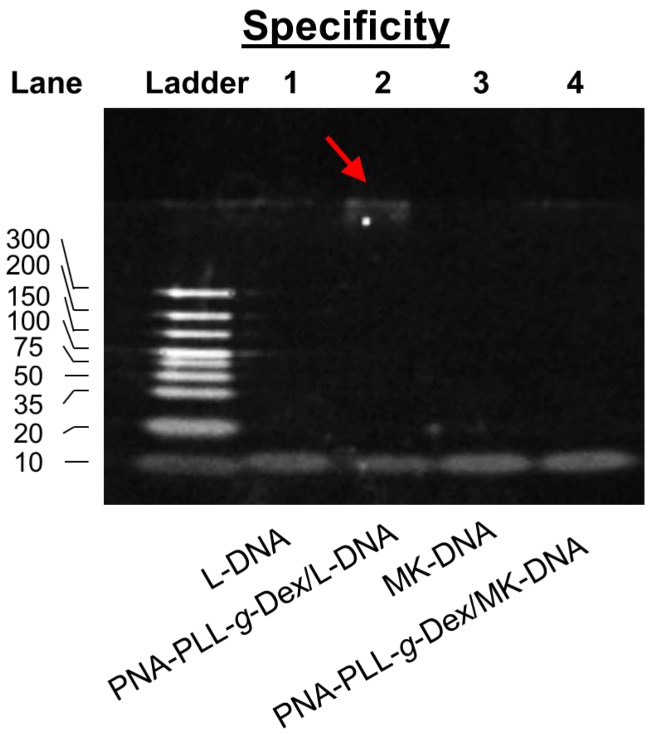


**Figure S2.** Specificity check of hybridization of PNA-PLL-g-Dex.

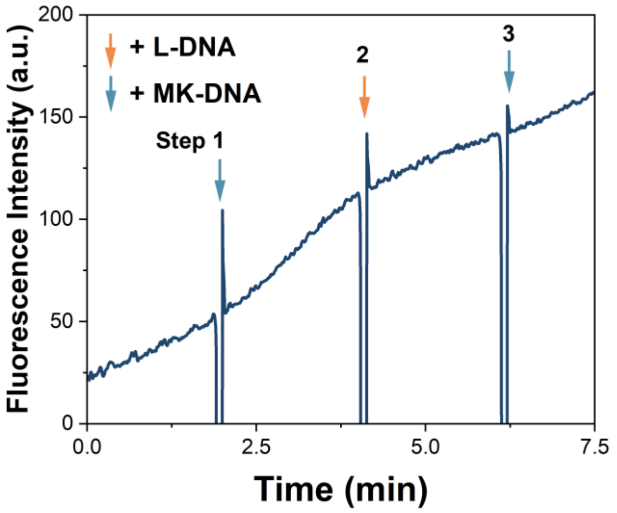


**Figure S3.** Zoomed range of fluorescence intensity of the MNAzyme system showing rapid switches of reaction upon repetitive addition of MK-DNA and L-DNA.

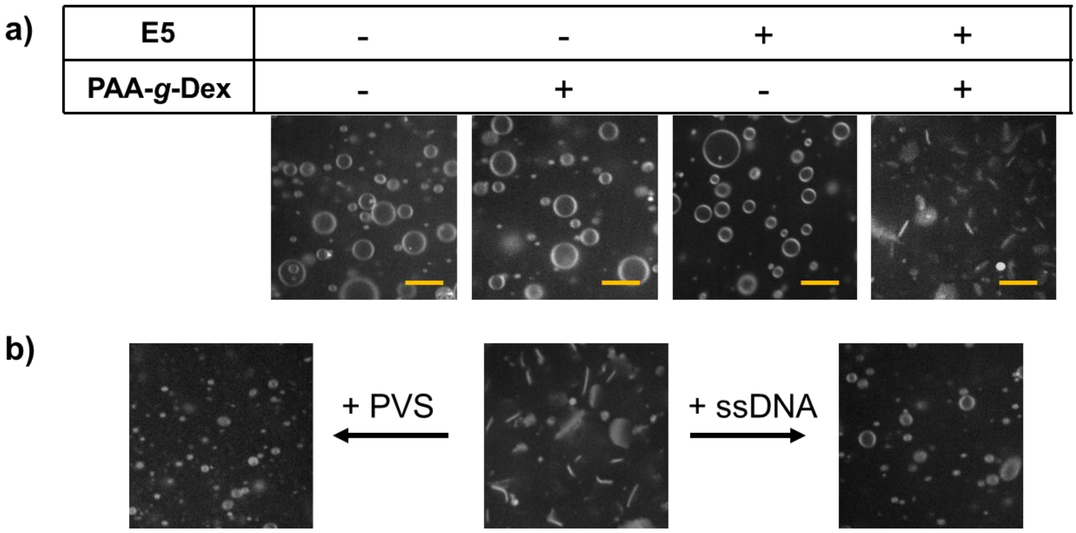


**Figure S4.** a) Vesicle-to-sheet conversion through E5 peptide and PAA-g-Dex under confocal laser scan microscopy (CLSM). b) Sheet-to-vesicle conversion driven by polyanions including poly(vinylsulfonic acid) (PVS) and single strand DNAs (ssDNA).

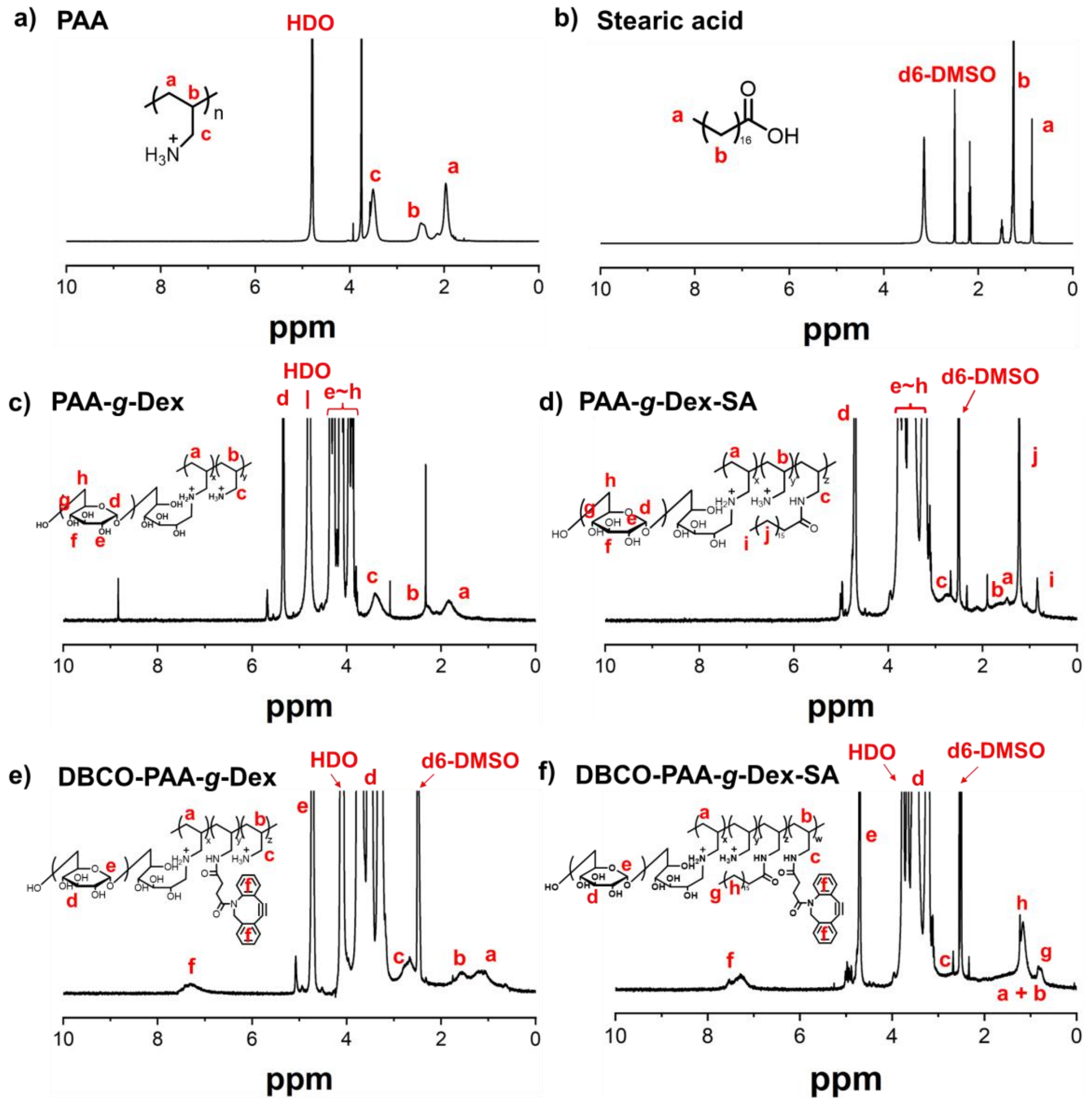


**Figure S5.** $^{1}$H-NMR spectra (400 MHz) of a) PAA, b) stearic acid [1], c) PAA-*g*-Dex, d) PAA-g-Dex-SA, e) DBCO-PAA-*g*-Dex and f) DBCO-PAA-*g*-Dex-SA, respectively.

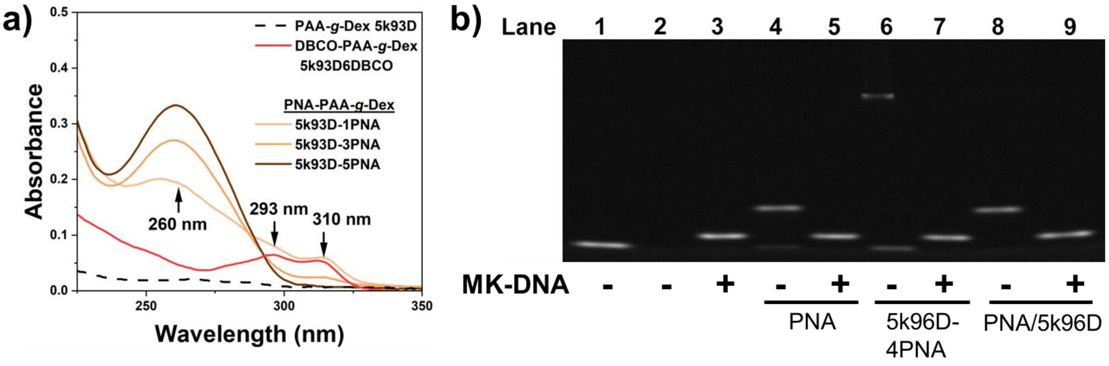


**Figure S6.** Synthesis of PNA-PAA-*g*-Dex. a) UV-Vis absorption of PAA-*g*-Dex 5k93D, DBCO-PAA-*g*-Dex 5k93D6DBCO and PNA copolymers of 5k93D-1PNA, 5k93D-3PNA and 5k93D-5PNA respectively in $H_2O$. b) Hybridization and toehold displacement performance of PNA-copolymer conjugates, in comparison to PNA-copolymer mixture and PNA without copolymers using native-PAGE electrophoresis (4.5% stacking gel and 20% separation gel). Lane 1: L-DNA; lane 2: PNA-PAA-g-Dex 5k96D-4PNA, lane 3: L-DNA/ MK-DNA hybrids; lane 4: PNA annealed with L-DNA before the addition of L-DNA and lane 5: after the addition of L-DNA; lane 6: 5k96D-4PNA annealed with L-DNA before the addition of L-DNA and lane 7: after the addition of L-DNA; lane 8: PNA annealed with L-DNA in presence of 5k96D before the addition of L-DNA and lane 9: after the addition of L-DNA.

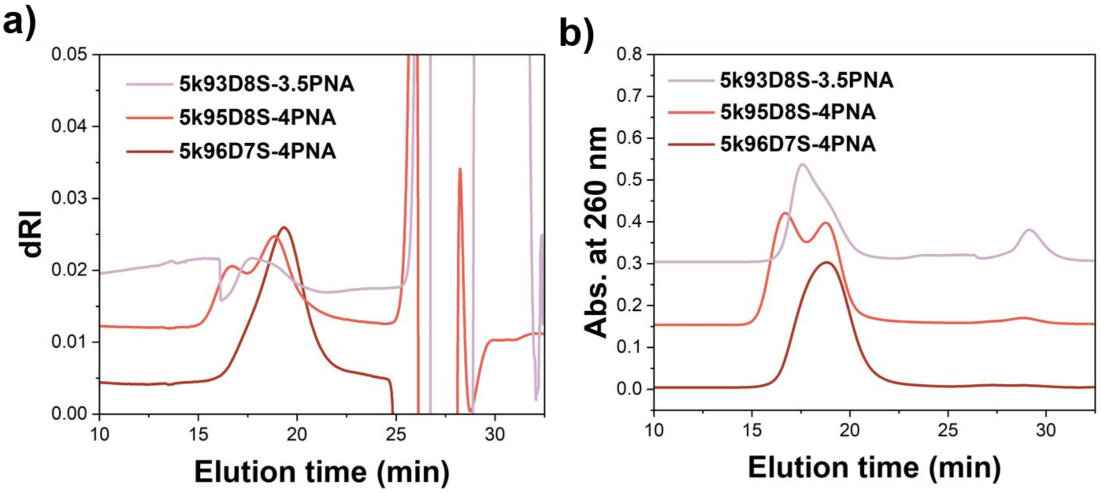


**Figure S7.** GPC profiles of PNA-PAA-g-Dex-SA with different dextran contents. a): dRI profiles, and b): UV absorption at 260 nm.

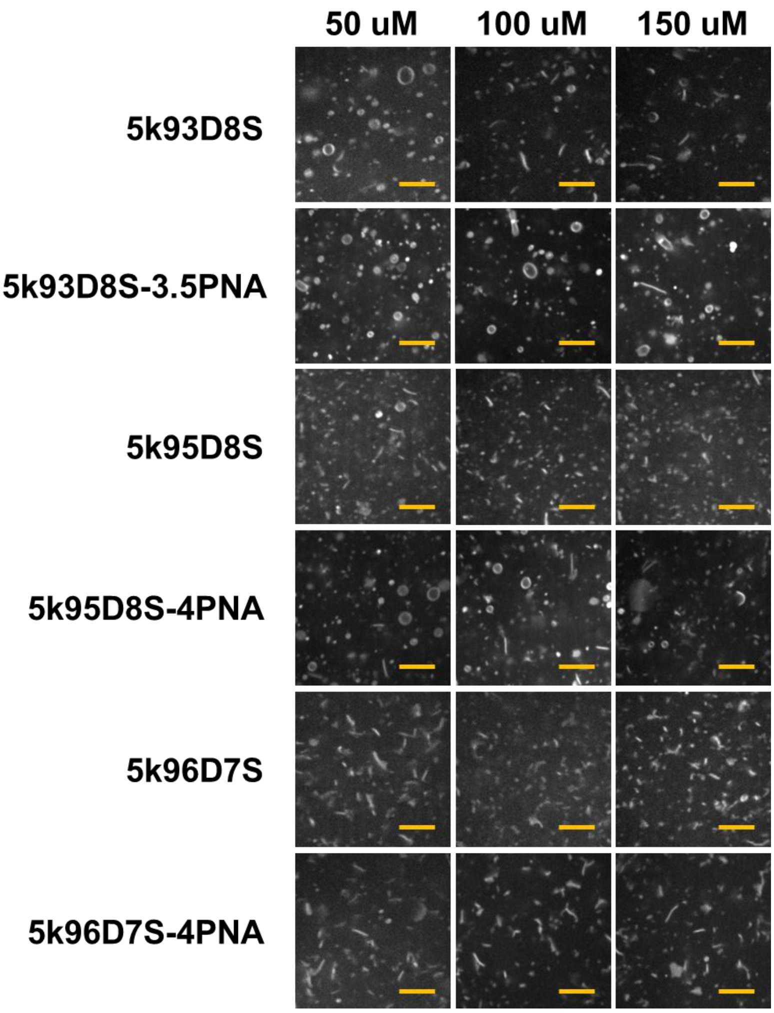


**Figure S8.** a) Assessment of nanosheets formation with PAA-g-Dex-SA and PNA-PAA-g-Dex-SA respectively with different copolymer concentrations, with the fixed lipid concentration of 500 µM and E5-pal of 7.5 µM using CLSM. Scale bar: 5 µm. b) The quantified sheet fractions of each copolymer from confocal microscopic images.

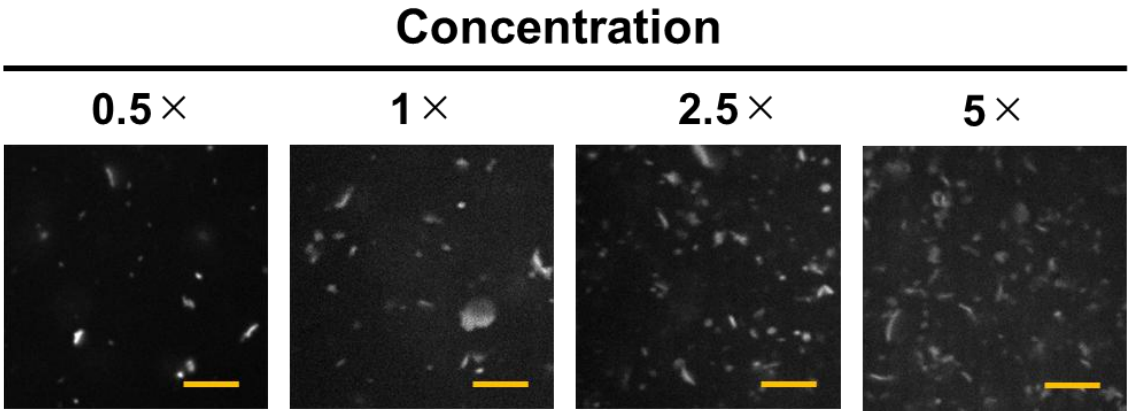


**Figure S9.** Confocal images of nanosheets formed with different folds of concentrations of lipids, 5k96D7S and E5-pal (1×: 60 µM lipids, 6 µM 5k96D7S and 0.9 µM E5-pal). Scale bar: 5 µm.

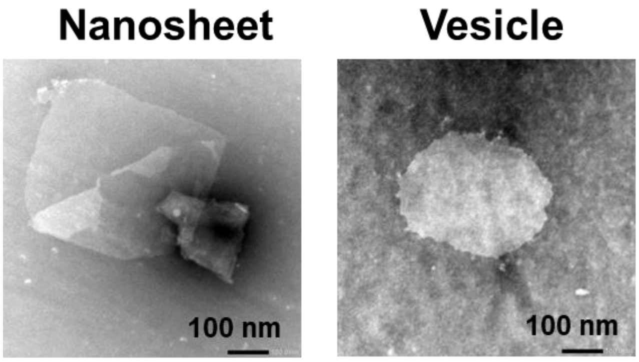


**Figure S10.** Typical morphology of nanosheet (left) and vesicle (right) observed by nsTEM, respectively. Magnitude: 50,000×.

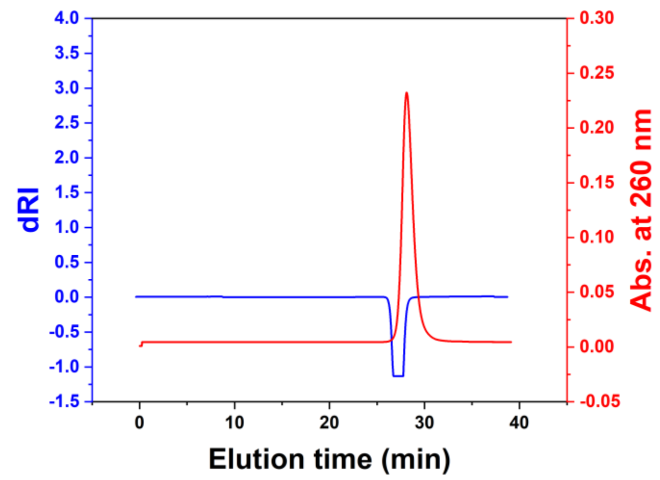


**Figure S11.** Typical GPC profiles of azide-PNA.
